# Microtubule dynamics is required for rapid coiling of haptonemata in haptophyte algae

**DOI:** 10.1101/322453

**Authors:** Mami Nomura, Kohei Atsuji, Keiko Hirose, Kogiku Shiba, Takeshi Nakayama, Ken-ichiro Ishida, Kazuo Inaba

**Author notes:** These two authors contributed equally to this work (Mami Nomura, Kohei Atsuji). Correspondence to Kazuo Inaba.

## Abstract

A haptonema is an elongated microtubule-based motile organelle uniquely present in haptophytes. The most notable and rapid movement of a haptonema is “coiling”, which occurs within a few milliseconds following mechanical stimulation in an unknown motor-independent mechanism. Here, we analyzed the coiling process in detail by high-speed filming and showed that haptonema coiling was initiated by left-handed twisting of the haptonema, followed by writhing to form a helix from the distal tip. On recovery from a mechanical stimulus, the helix slowly uncoiled from the proximal region. Electron microscopy showed that the seven microtubules in a haptonema were arranged mostly in parallel but that one of the microtubules often wound around the others in the extended state. The persistence lengths calculated from the curvature of the haptonematal microtubules indicated their unusual flexibility. A microtubule stabilizer, paclitaxel, inhibited coiling and induced right-handed twisting of the haptonema in the absence of Ca^2+^, suggesting changes in the microtubule surface lattice. Addition of Ca^2+^ caused bend propagation toward the proximal region. These results indicate that switching microtubule conformation with the aid of Ca^2+^-binding microtubule-associated proteins is responsible for rapid haptonematal coiling.

**Summary Statement:** Microscopy observations and pharmacological experiments revealed that the rapid coiling of a non-motor microtubule-based motile organelle, the haptonema, is explained by conformational changes of microtubules, including twisting and writhing.

## INTRODUCTION

Haptophytes are the group of microalgae that are widely distributed in oceans. They show similarities to heterokonts in chloroplast structure and chlorophyll species but are classified into an independent phylum owing to several cytological properties, including the lack of mastigonemes on flagella and the presence of extracellular scales or coccoliths (Christensen et al., 1962; Andersen, 2004). A haptonema is a filiform organelle uniquely present in haptophytes (Parke et al., 1955). It extends from a position between the bases of two flagella, reaching up to more than 100 μm in some species (Gregson et al., 1993a,b). A variety of functions have been demonstrated for haptonema, including attachment and gliding on a substrate, formation of food aggregates, food capture and transport, and reception of mechanical stimuli (Manton, 1967; Leadbeater and Manton, 1971; Kawachi and Inoue, 1991, 1995). When haptophytes receive mechanical stimuli, they fully coil the haptonema within only a few milliseconds. By contrast, “uncoiling”, the process to resume the extended state, is much slower than coiling. The coiling state is thought to be a low-energy form, because the haptonema is always coiled when it is detached or when haptophytes are dead (Estep and Maclntyre, 1989).

Haptonema coiling is inhibited by EGTA depletion of extracellular Ca^2+^ or by the addition of an inhibitor of Ca^2+^-induced Ca^2+^ release, indicating that it is triggered by Ca^2+^ influx followed by efflux from a Ca^2+^ store (Kawachi and Inouye, 1994). Upon receipt of a mechanical stimulus, haptonematal coiling accompanies a change in waveform and an increase in beat frequency of flagella, possibly from changes in intracellular Ca^2+^ concentrations. This results in a quick response to avoid the stimulus (Gregson et al., 1993b; Kawachi and Inouye, 1994). The mechanism for the cytoskeletal response to Ca^2+^ is not well understood, except that centrin is localized as a small dot-like structure at the distal tip of haptonematal microtubules (Lechtreck, 2004).

Ultrastructural observations show that six to seven microtubules pass in parallel through a haptonema. In cross section, they are arranged in a circle of ~100 nm in diameter in the middle region of the haptonema, and in an arc-shape with invagination of endoplasmic reticulum at the basal region. The number of microtubules reduces to three at the distal most region (Manton, 1964, 1967, 1968; Gregson et al., 1993b). It is well known that microtubules of flagellar axonemes have post-translational modifications (Wloga and Gaertig, 2010), but the patterns of modifications of haptonema microtubules are different from those of axonemes (Lechtreck, 2004). The microtubules are surrounded by fenestrated cisternae in the major part of a haptonema beneath the plasma membrane. From thin-section electron microscopy observations, the circular arrangement of microtubules tends to change to a crescent arrangement after coiling. An electron dense structure in the center of the microtubule ring and a structure that links neighboring microtubules are observed (Gregson et al., 1993b). However, no structure that potentially represents motor proteins, such as dyneins or kinesins, has been observed. Thus, the molecular mechanism for rapid coiling of haptonemata is completely unknown.

Here we examined the structure of haptonemata and the process of their coiling using a newly identified marine species of the genus *Chrysochromulina*. Although the structures of this haptonema share common properties to those reported for other species (Gregson et al., 1993), we obtained new information regarding microtubule configurations in this haptonema. Furthermore, we found that a microtubule stabilizer, paclitaxel (taxol), inhibit s haptonematal coiling, indicating that microtubule flexibility and dynamics are responsible for the induction of rapid haptonema coiling.

## RESULTS

### Morphological characterization of *Chrysochromulina sp.* NIES-4122

The genus *Chrysochromulina* (Prymnesiophyceae) is characterized by the development of a relatively long haptonema in the subclass Chrysochromulinaceae (Edvardsen et al., 2011). Here, we used a *Chrysochromulina* species that was collected in Tokyo bay in 2013. The cell strain was established by clonal culture. This species has no calcareous coccoliths but has organic scales with no spine (Fig. S1A-C) and is morphologically classified into the genus *Chrysochromulina*. However, the shapes of the scales are distinct from any known *Chrysochromulina* species. This species possesses a haptonema of up to ~150 μm in length, which is a little longer than that in *C. simplex, C. acantha* and *C. hirta* (Fig. S1B; Kawachi and Inouye, 1995; *C. hirta* is changed to *Haptolina hirta* after Edvardsen et al., 2011). When compared with other *Chrysochromulina* species, such as NIES-1333, the haptonema of this species was more resistant to mechanical stimuli that cause detachment from the cell body. Coiling was partially inhibited by depletion of Ca^2+^ in artificial sea water and completely inhibited by chelating intracellular Ca^2+^ (Fig. S1D). This strain has been deposited with the National Institute for Environmental Studies (NIES), Japan, as *Chrysochromulina* sp. NIES-4122.

### Observation of haptonematal coiling by high-speed recording

The haptonema of *Chrysochromulina* sp. NIES-4122 occasionally showed coiling during observation under a light microscope. Gentle tapping of the microscope stage induced almost 100% haptonematal coiling (Movie 1). The coiling occurred very rapidly and was complete within 5-10 msec (Movie 2), which is considerably faster than that observed in *C. acantha* (10-20 msec; Leadbeater and Manton, 1971). In contrast, uncoiling was much slower, taking ~480 msec to complete extension (Movie 3).

One might expect that the coiling would start from the tip of a haptonema. However, detailed observation of the high-speed images revealed that this is not the case; the distal half of the haptonema first began to bend in gentle helices, followed by sequential coiling from the tip (Fig. 1A). The coil appeared to be left-handed, which was more clearly observed in the process of uncoiling (Fig. 1B). Uncoiling initiated from the proximal region of a haptonema while the distal most part remained curled, which was then gradually unwound during the last step of extension.

**Fig. 1.**
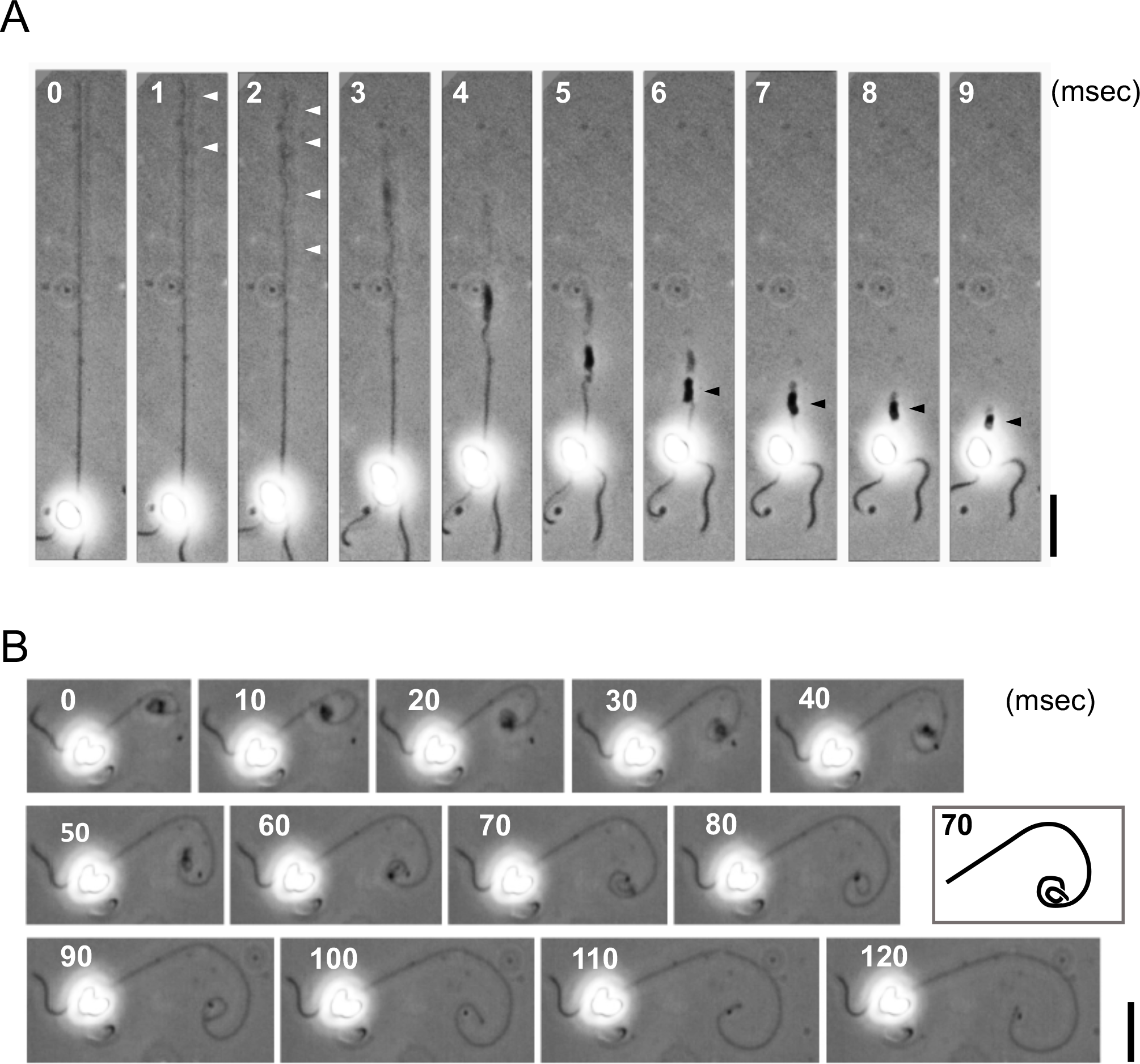
High-speed analysis of haptonematal coiling and uncoiling. **(A)** Coiling process of a haptonema. High speed images recorded at 1000 fps. Gentle helices that initially formed in the distal half of a haptonema are indicated by white arrowheads. Stacked coils are indicated by black arrowheads. Bar, 20 μm. **(B)** Uncoiling process of a haptonema. High speed images recorded at 200 fps. Bar, 20 μm. The inset represents the trace of the haptonema at 70 msec for clarification of the coiling direction.

### Microtubule configuration in haptonemata in extended and coiled states

As reported in other species, including *C. chiton* (Manton, 1967), *C. simplex* and *C. acantha* (Gregson et al., 1993b), thin-section electron microscopy showed that in the extended haptonema seven microtubules are arranged in a ring, which is peripherally surrounded by cisternae (Fig. 2A-D). This circular arrangement of microtubules was distorted in the coiled haptonema and one of the microtubules was often invaginated towards the center (Fig. 2E-H). We measured the center-to-center distances between adjacent microtubules (Fig. S2A, B). The distances were relatively constant in an extended haptonema among sequential sections (Fig. S2C) but in a coiled haptonema one of the inter-filament distances sometimes became deviated, as if the ring was torn open (Fig. S2D). The deviated microtubule often changed its position relative to the adjacent microtubule (Fig. S2D). This pattern with an interfilament deviation distance of more than 15 nm was observed in 0% and 13% of extended and coiled haptonemata, respectively.

**Fig. 2.**
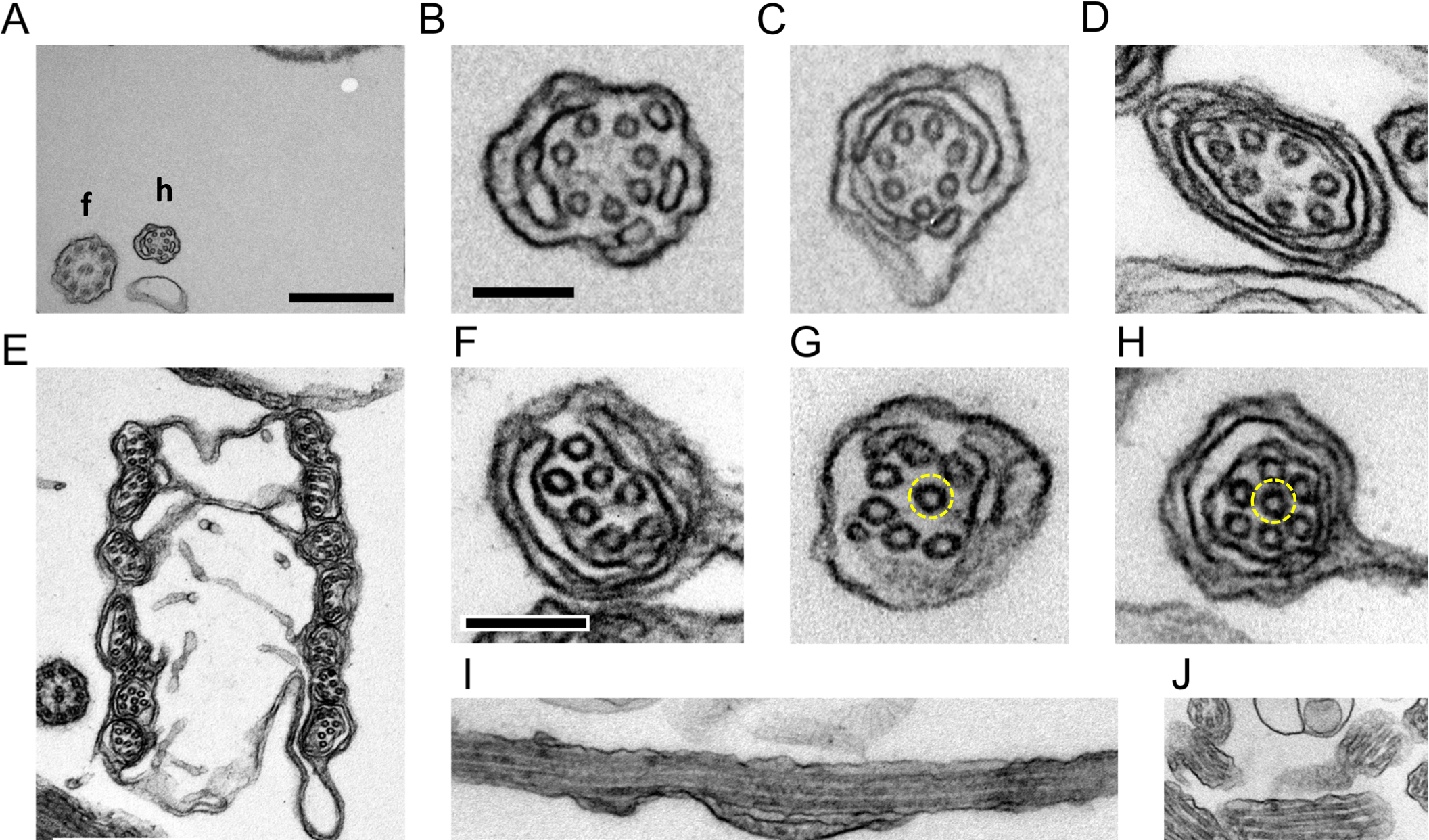
Thin-section images of extended and coiled haptonemata. **(A)** Extended haptonema at low magnification. Bar, 500 nm. (**B-D**) High magnification images of an extended haptonema. Bar, 100 nm. **(E)** Coiled haptonema at low magnification. Bar, 500 nm. (**F-H**) High magnification images of a coiled haptonema. Dashed yellow circles show a microtubule invaginated into the center. Bar, 100 nm. (**I and J**) Longitudinal images of extended (I) and coiled (J) haptonemata. Bar, 500 nm.

In longitudinal sections 70–80 nm thick, we were able to observe a parallel arrangement of three, sometimes four microtubules of up to 1 μm and 200 nm in extended and coiled haptonemata, respectively (Fig. 2I, J). This indicates that microtubules are arranged in a more or less parallel manner in both extended and coiled stages. To confirm the parallel arrangement of microtubules, we treated a small plate of polymerized Epon resin with poly-lysine. This was then coated with bovine serum albumin. Fixed haptophytes were deposited on the coated resin, fixed and embedded in Epon. This procedure provided a fixed landmark to measure the relative positions of each microtubule between sequential sections (Fig. 3A-C). The position of each microtubule relative to a landmark would change periodically if the haptonematal microtubules were arranged helically. In the case of parallel arrangement, the positions of seven microtubules would shift in parallel in the sequential sections (Fig. 3D). These results indicated that the microtubules are arranged in parallel, at least up to lengths of 400 nm and 320 nm for the extended and most of the coiled haptonemata, respectively, (Fig. 3E, F). However, four out of 24 sequential images of the coiled haptonema showed twisted arrangements (Fig. 3G).

**Fig. 3.**
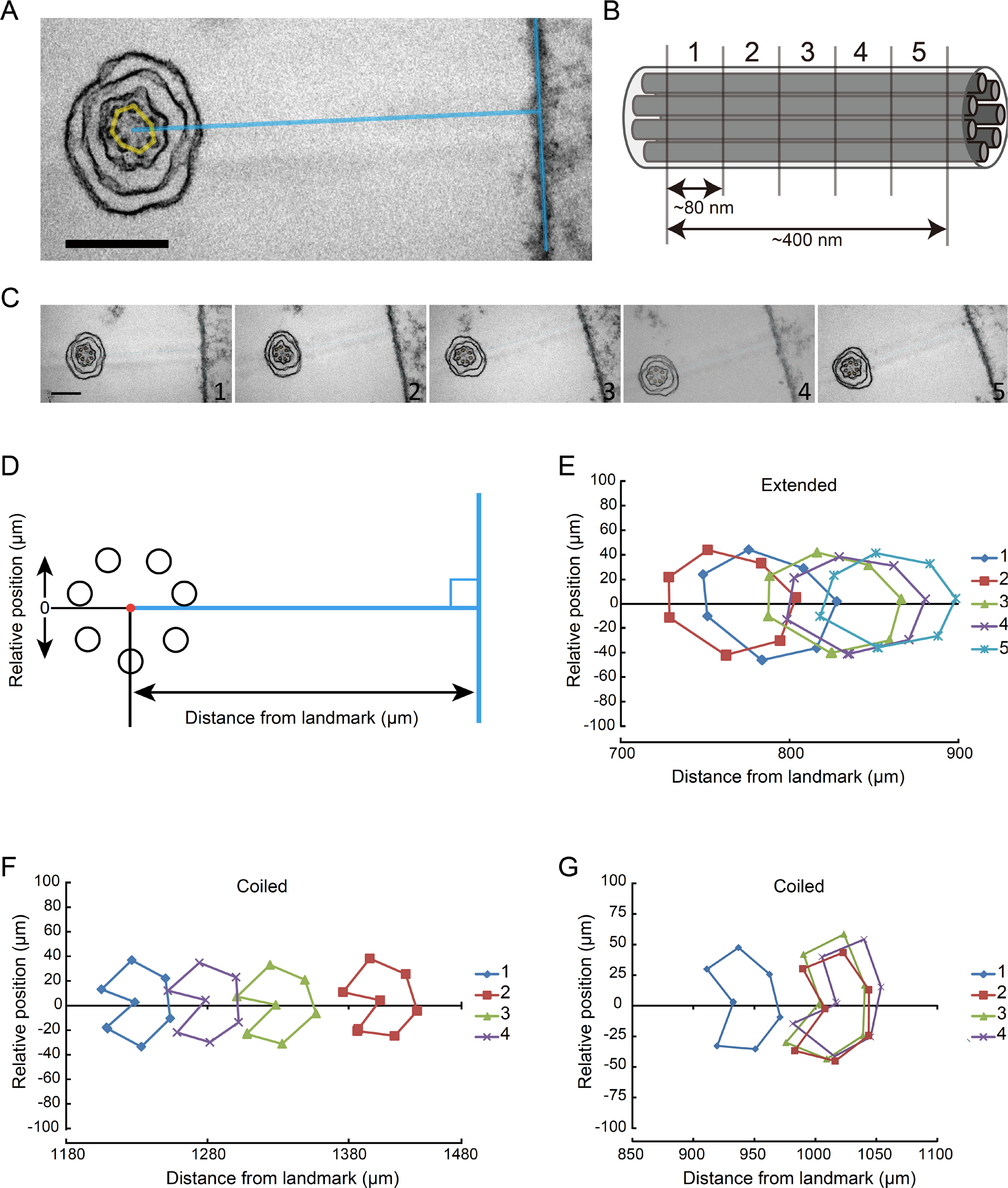
Arrangement of microtubules along a haptonema. **(A)** Extended haptonema adjacent to the poly-lysine-BSA coated Epon surface (landmark). To position each microtubule, a line was drawn from the center of the microtubule bundle at right angles to the surface. Bar, 200 nm. (**B**) Approximately 80 nm sequential sections were made. For example, five sections covered a 400 nm length of a haptonema along the longitudinal axis. (**C**) Example of sequential images of an extended haptonema. Bar, 200 nm. (**D**) Definition of the distance from the landmark and the relative position of each microtubule. (**E**) Typical plot of microtubule positions in five sequential images of an extended haptonema. (**F**) Typical plot of microtubule positions in five sequential images of a coiled haptonema. (**G**) Plot showing a helical arrangement of the seven microtubules in a coiled haptonema. Four out of 24 sets of sequential images of the coiled haptonema showed this pattern.

Next, haptophytes were deposited on a grid, demembranated by NP-40 and observed by negative staining electron microscopy (Fig. 4). The diameters of flagellar axonemes and haptonemata were distinct, so that a long haptonema could be distinctly observed at lower magnifications by negative staining (Fig. 4A). As reported by Gregson et al. (1993b), haptonematal microtubules are bound together after demembranation but are partly dissociated in some regions of the extended haptonema. Microtubules of the extended and coiled parts of a haptonema are mostly parallel to each other without any torsion. However, we often observed a microtubule loosely wound around the other six microtubules (Fig. 4B). This peculiar microtubule winding was not clearly observed in the coiled region where all the microtubules are mostly arranged in parallel. Instead, we observed that microtubules were stacked at two opposite positions in the coil (Fig. 4C). This is compatible with the observations from thin-sectioned sequential images of some microtubules being twisted in the coiled region (Fig. 3).

**Fig. 4.**
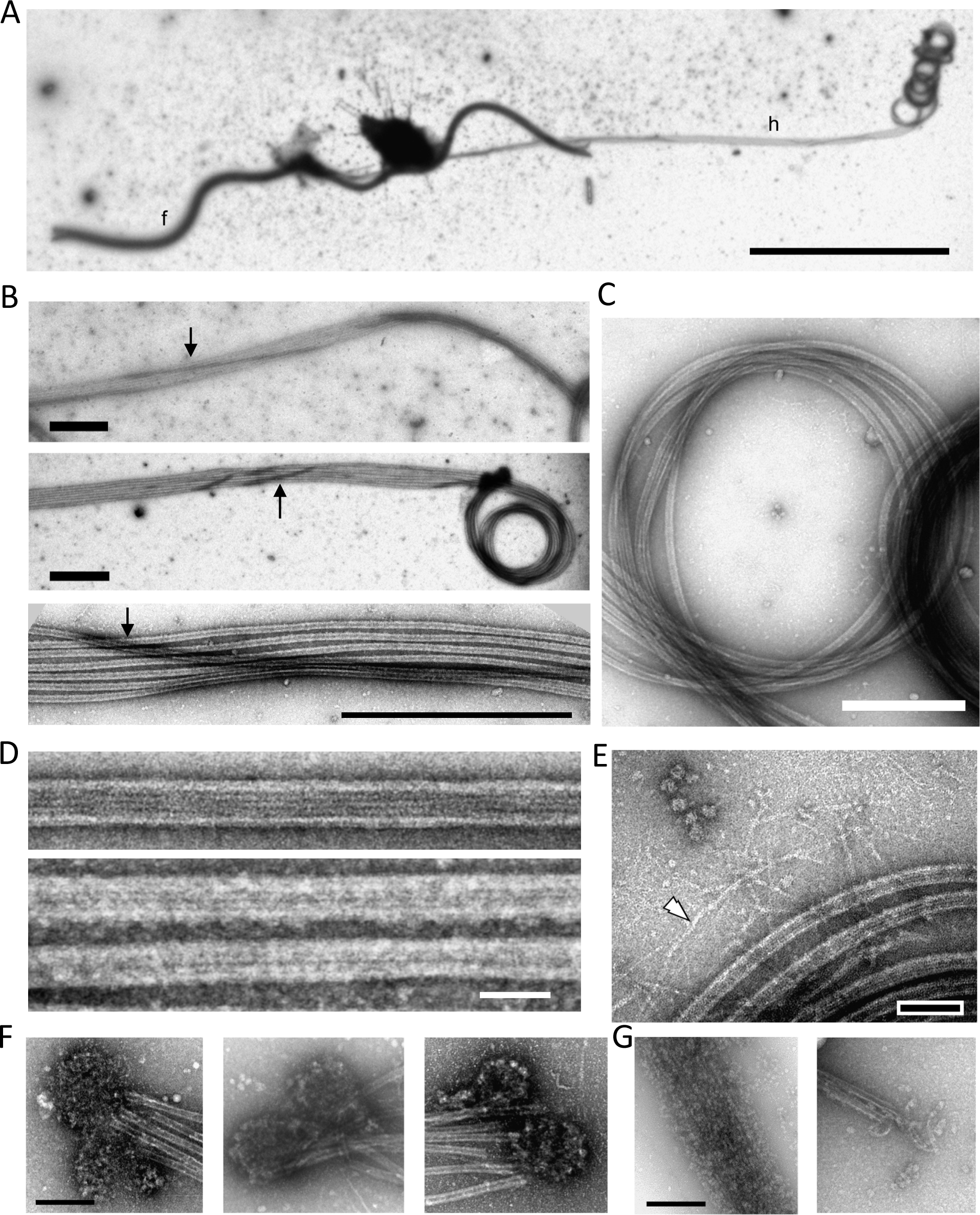
Negative stain images of extended and coiled haptonemata. **(A)** Negative stain image of a haptophyte. Because of demembranation, most parts of the cell body are removed. f, flagellum; h, haptonema. A long haptonema is observed on the right side with the tip coiled. Bar, 10 μm. (**B**) Three images showing that one, occasionally two microtubules (arrows), wind around the other microtubules in the extended region. Bar, 1 μm. (**C**) Coiled region. Microtubules are mostly parallel to each other, but there seems to be a twist at the two opposite regions of the coil (arrowhead). Bar, 500 nm. (**D**) Microtubules polymerized from purified brain tubulin (upper) and microtubules in a demembranated haptonema (lower). Bar, 50 nm. (**E**) Image showing filamentous structures (white arrowhead) emanating from the microtubule bundle. Bar, 100 nm. (**F**) Mass of small particles observed at the distal tip of a haptonema. They are frequently observed in a pair. Bar, 200 nm. (**G**) Depolymerization of microtubules occasionally observed in demembranated haptonemata. The depolymerized microtubules are still bundled probably because of MAPs. The morphology of the small particles in *F* is similar to depolymerized tubulin dimers. Bar, 200 nm.

The diameter of the haptonema coil was 1.2 μm ± 0.03 μm (N = 83), as estimated from negative stain images. The microtubules in the coils are largely curved (Fig. 4C), suggesting higher flexibility than microtubules from other sources. To evaluate the flexibility of haptonematal microtubules, we calculated the persistence length, a measure of flexural rigidity of filamentous structures. The persistence lengths for DNA, an actin filament, and a microtubule have been reported to be 50 nm, 13 μm, and ~6 mm, respectively (Taylor and Hagerman, 1990; Gittes et al., 1993; Käs et al., 1994), indicating that microtubules are very rigid structures. However, the persistence lengths of haptonematal microtubules calculated according to the method of Gittes et al. (1993) for the extended and coiled regions of the haptonema were 93.1 ± 17.7 μm (N = 57) and 1.7 μm ± 0.2 μm (N = 78), respectively (Fig. S3). Although these values were calculated using the negative stain images of microtubules fixed on a carbon membrane and thus do not represent the true values of the persistence length, the results indicate that haptonematal microtubules have high flexibility.

It is likely that binding of haptonema-specific microtubule-binding proteins (MAPs) prevents the curved microtubules from depolymerization. In fact, negative stain images comparing haptonematal microtubules (Fig. 4D, lower) and microtubules polymerized from purified tubulin (Fig. 4D, upper) showed that the haptonematal microtubules are covered with additional proteins. When haptonematal microtubules were depolymerized, they often appeared to stay bundled (Fig. 4G, left), again indicating the presence of large amounts of MAPs. We also occasionally observed filamentous structures with diameters that were much smaller than those of microtubules. They were observed spreading out from microtubule bundles (Fig. 4E). Another unique structure we observed was a mass of small particles. These masses were observed at the distal tip of a haptonema and usually existed as a pair (Fig. 4F), suggesting a cap structure at the distal end of haptonematal microtubules. Alternatively, they may simply be masses of tubulins depolymerized from the distal part of haptonematal microtubules because they are structurally similar to the depolymerized microtubules (Fig. 4G).

### Taxol inhibits haptonematal coiling

Ultrastructural observation of the haptonema indicated dynamic structural changes between coiled and extended states. To examine the possibility that microtubule dynamics are involved in the coiling mechanism, we used taxol and nocodazole, which inhibit microtubule depolymerization and polymerization, respectively. We incubated the haptophytes with each drug and induced coiling by tapping the microscope stage. Nearly 80% of nocodazole-treated and control haptophytes showed coiling. However, haptophytes treated with taxol showed significantly inhibited coiling (Fig. 5A). In the extended state without mechanical stimulation, control haptonemata remained straight but taxol-treated haptonemata showed bending with low curvatures. The overall length of taxol-treated haptonema was significantly increased (Fig. 5B), which might indicate elongation of the microtubules, possibly from the tubulin pool at the distal tip. Decreased length was observed for nocodazole-treated haptonema but this was not significant.

**Fig. 5.**
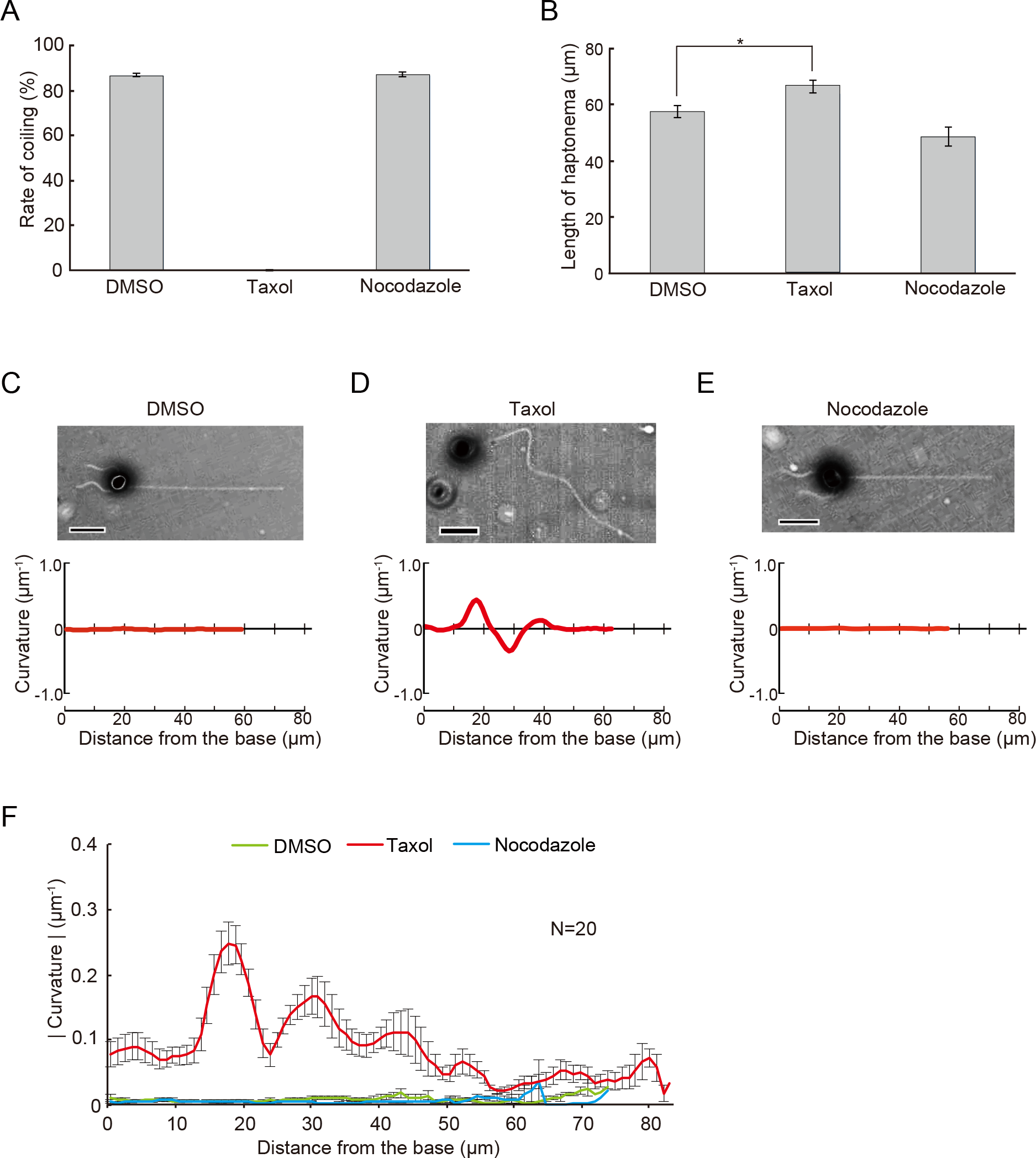
Effects of drugs that affect microtubules on haptonematal coiling. **(A)** Rates of coiling in the presence of taxol or nocodazole. Error bars show the standard deviation. N=5. (**B**) Length of haptonemata at 60 min after treatment with each drug. Error bars show the standard deviation. N=5. The asterisk represents that the difference is significant at p < 0.01 (Student’s t-test). (**C-E**) Effects of each drug on the curvature along the axis of a haptonema. Corresponding phase-contrast images are shown above each plot. Bar, 10 μm. (**F**) Absolute values of the curvature showing the extent of bending along the haptonema. Error bars show the standard deviation. N=20.

To quantitatively evaluate the taxol-induced bending, we measured the curvature along the haptonema (Fig. S4). The curvature was almost zero in control and nocodazole-treated haptophytes throughout the haptonema (Fig. 5C, E), but taxol-treated haptonema showed a large peak in the proximal region with the maximum at around 15-20 μm from the base (Fig. 5D, F; Fig. S5). This bending was initially formed in the distal to middle region 5 min after treatment and then propagated toward the proximal region (Fig. S6A).

### Changes of taxol-induced haptonema bending by Ca^2+^

To explore the relationship between the mechanism of coiling and taxol-induced bend formation, we examined the effect of Ca^2+^ on haptonemata after treatment with taxol in Ca^2+^-free conditions. In contrast to taxol-treated haptonemata in normal sea water with Ca^2+^ (Fig. 5D), the haptonema in Ca^2+^-free conditions did not show planar bending but rather twisted or helical shapes (Fig. 6A). Careful observation by altering the microscope focus showed that the helix was right-handed. The helix was maintained from 5 min to 60 min but became gentle in pitch to extend toward the tip (Fig. S6B).

**Fig. 6.**
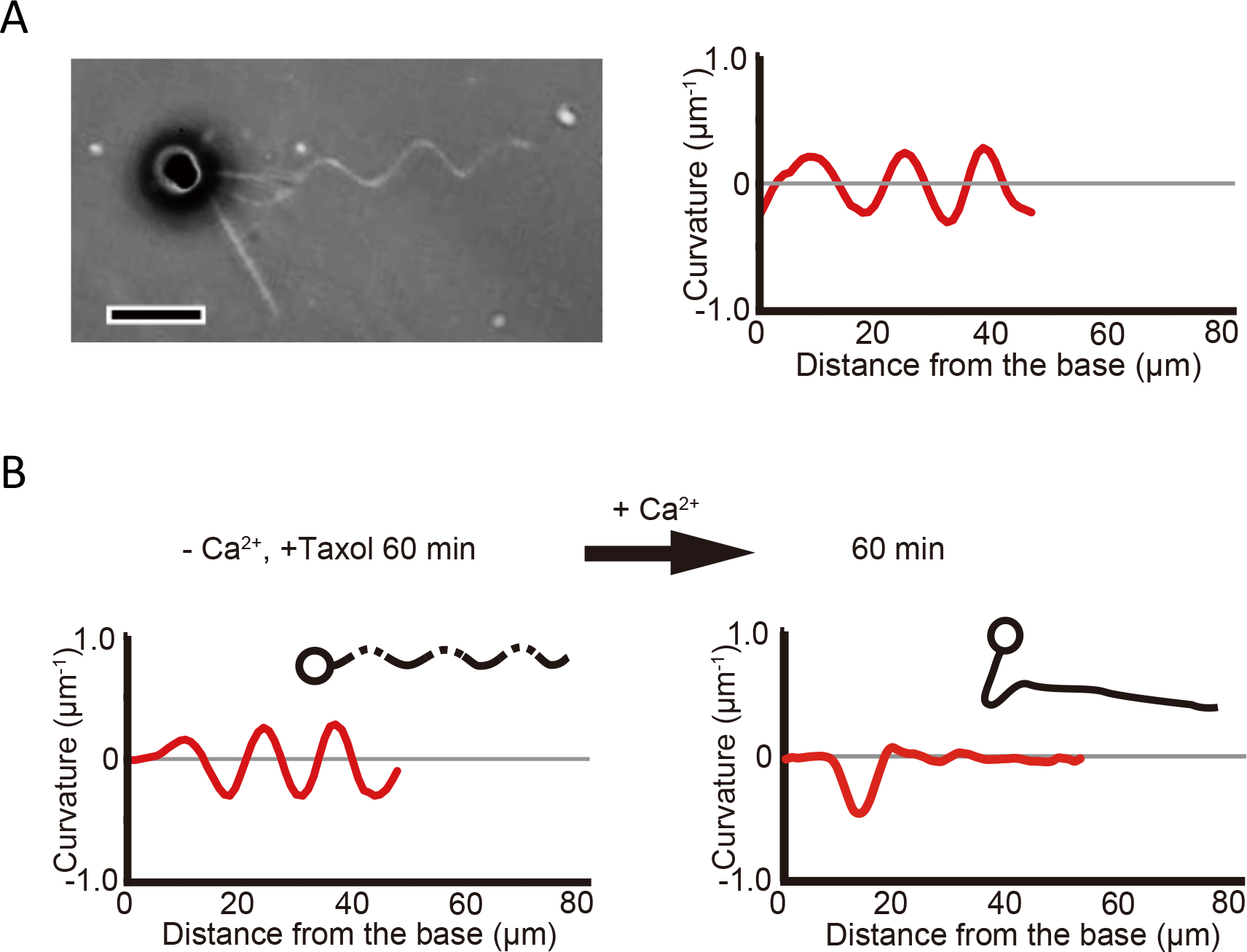
Effects of Ca^2+^ on taxol-induced haptonematal bending. **(A)** A phase contrast image of a haptophyte treated with taxol in the absence of Ca^2+^ (CFSW+EGTA+BAPTA-AM) showing a helical configuration of the haptonema (left). The plot to the right shows the curvature along the haptonema. Bar, 10 μm. (**B**) Ca^2+^-dependent propagation of a taxol-induced haptonematal helix. The helix is also converted to planar bending with the maximum curvature of bend propagating to the proximal region. Trace images of haptophytes are shown as insets.

To examine the requirement of Ca^2+^, taxol-treated haptonemata were first incubated in Ca^2+^-free conditions containing EGTA/BAPTA-AM for 60 min, and then 50 mM CaCl_2_ was added. The twist observed 60 min after the incubation in Ca^2+^-free conditions gradually became planar and shifted toward the base of the haptonema after the addition of CaCl_2_ (Fig. 6B). The final waveform became similar to that observed in taxol-containing artificial sea water (Fig. 5D).

## DISCUSSION

Our present data demonstrate that the quick coiling of haptonema is driven by the configurational changes and dynamics of seven singlet microtubules, not through microtubule-sliding driven by conventional molecular motors. Another eukaryotic example showing non-motor rapid movement by microtubule structures is the contraction of heliozoan axopodia (Tilney and Porter, 1965; MacDonald and Kitching, 1967; Suzaki et al., 1980; Hausmann et al., 1983), in which Ca^2+^-dependent cataclysmic breakdown of microtubules as well as their depolymerization induce the contraction (Febvre-Chevalier and Febvre, 1986). However, heliozoan axopodia do not make coiled structures, and our data showed that the microtubules in coiled haptonemata are not depolymerized. An example showing a similar movement to haptonematal coiling is quick contraction of the spasmoneme in the “stalk” of the peritrichous ciliates, such as *Vorticella* and *Zoothamnium*. The spasmoneme contraction proceeds very quickly to form a spiral stalk in a Ca^2+^-dependent but ATP-independent manner. Cytoskeletal elements of the stalks are not microtubules but helically coiled structures, mainly constructed of a 20 kDa centrin-related Ca^2+^-binding protein, called spasmin (Amos et al., 1975). Thus, haptonematal coiling is a unique type of microtubule-dependent motility, distinct from any other known eukaryotic motile machinery.

### Microtubule dynamics is involved in haptonematal coiling

We found that taxol inhibited the rapid coiling of a haptonema. Although this inhibition may indicate involvement of microtubule depolymerization in the coiling process, electron microscopy showed no depolymerization of microtubules in the coiled haptonema. Thus, the inhibition is considered to be caused by another mechanism. Taxol binds to β-tubulin, resulting in a longitudinal extension of the αβ-tubulin dimer spacing, formation of straight protofilaments and stabilization of microtubules (Arnal and Wade, 1995; Caille et al., 2007; Mitra and Sept, 2008). Application of taxol to polymerized microtubules significantly changes their flexural rigidity (Dye et al., 1993; Mickey and Howard, 1994; Venier et al., 1994) and overall radial flexibility (Donhauser et al., 2010). Therefore, the gentle coiling of haptonema in the presence of taxol is likely to be caused by changes in the mechanical property of microtubules.

The flexural rigidity of biological filaments is often evaluated using ‘persistence length’. Compared with double-stranded DNA or actin-filaments, whose persistent lengths are reported to be 50 nm (Taylor and Hagerman, 1990) and 10 μm (Gittes et al., 1993; Käs et al., 1994), respectively, microtubules are rigid filaments and their persistence length is large, ranging from 80 μm to 5.2 mm depending on the measurement condition (Gittes et al., 1993; van den Heuvel et al., 2008). Persistence lengths also depend on filament length, and microtubules covalently grafted to a grid showed persistence lengths ranging from 110 to 5,035 μm for filament lengths of 2.6 to 47.5 μm (Pampaloni et al, 2006). We obtained persistence lengths of 93.1 and 1.7 μm in the extended and coiled regions of haptonemata, which are the smallest values among those reported for microtubules. Notably, the microtubules in coiled haptonemata showed unusually curved structures, which cannot be explained by previously-reported persistence length values. Such a high curvature is expected to cause a large distortion of the microtubule surface lattice.

Formation of helical shapes (Fig. 6A) by taxol treatment should be accompanied by distortion of the tubulin lattice. Taxol alters the tubulin lattice and supertwist of microtubules (Arnal and Wade, 1995); therefore, it is possible that the microtubule structures of taxol-treated haptonemata in the absence of Ca^2+^ are distorted and fixed to make the right-handed helical shape. When trapped in this structural state, the changes in configuration needed for the left-handed coiling of haptonemata may be inhibited. It is likely that haptonematal microtubules need extra structural reinforcement to prevent them from breakage during coiling (Fig. 4D). The microtubule destabilizer, nocodazole, induces no apparent morphological change in haptonemata, such as shortening by depolymerization, indicating the presence of structural reinforcement for microtubule stability. Together, these data indicate that the coiling of haptonematal microtubules is caused by mechanical changes of microtubules with the aid of specific MAPs.

### The role of Ca^2+^ in configuration changes of haptonematal microtubules

Haptonemata coiling occurs within 5–10 msec; therefore, haptonematal microtubules are thought to have a structural basis that allows a rapid change in conformation in response to an increase in intracellular Ca^2+^. Since *in vitro* polymerized microtubules are straight, a mechanical strain must be imposed on microtubules for coiling. This strain is probably induced by MAPs, among which Ca^2+^-binding proteins seem to play major roles in keeping distinct microtubule configurations in the presence of Ca^2+^ (Fig. 7). Ca^2+^ binding to the haptonematal MAPs alters their conformations, imposing a strain on the microtubule (Fig. 7B). This strain may be relaxed when the microtubule coils (Fig. 7C). The haptonemata of dead cells and detached haptonemata are always coiled (Leadbeater and Manton, 1969; Gregson et al., 1993b), and the intracellular Ca^2+^ of these haptonemata is high. Therefore, the coiled configuration of haptonematal microtubules seems to be stable in the presence of Ca^2+^-bound MAPs. When Ca^2+^ is released from the MAPs bound to the coiled microtubule, the microtubule conformations change back to the extended form (Fig. 7D). The conformation of MAPs is assumed to be different between Ca^2+^-bound (Fig. 7B) and Ca^2+^-free (Fig. 7D) forms. This might result in the different microtubule configurations observed during coiling (gentle helices) and uncoiling (uncurling) (Fig. 1).

**Fig. 7.**
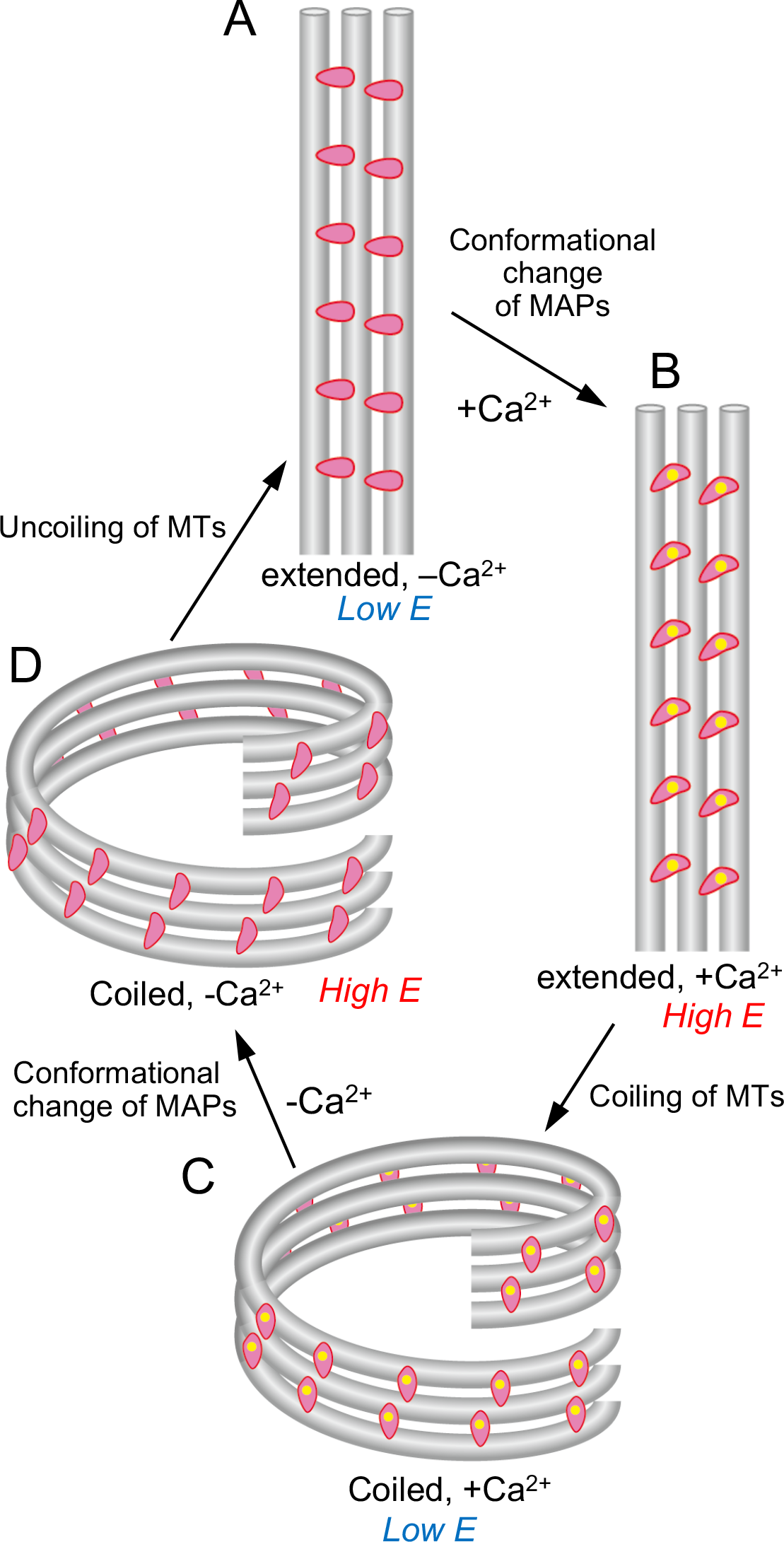
Model for configuration changes in haptonematal microtubules driven by Ca^2+^-binding MAPs. The three grey tubes represent either the three neighboring microtubules in a haptonema or three neighboring protofilaments in a haptonematal microtubule. In the former case, the microtubules are linked by Ca^2+^-binding MAPs (pink). When Ca^2+^ is not bound to the MAPs, the microtubules are stable in the straight conformation (A). When Ca^2+^ (yellow) binds, the MAPs change their structures, causing a shearing force between the microtubules (B). The tension is reduced when the microtubule bundle becomes coiled (C). When Ca^2+^ is released from the MAPs, their structure changes to a conformation different from the Ca^2+^-bound form in B (D). This conformational change induces uncoiling of the microtubules and the haptonema extends. In the case when the grey tubes represent protofilaments, the MAPs are bound to each protofilament. Ca^2+^-binding to the MAPs changes the structure of the protofilament and thus the interactions between neighboring protofilaments, which results in a torsional force exerted on individual microtubules (B). The force is relaxed when the microtubule becomes coiled (C). When Ca^2+^ is released, the MAPs take another conformation (D), which changes the inter-protofilament interactions, resulting in uncoiling. In either case, release of Ca^2+^ from the MAPs brings the microtubule conformation back to the extended state.

There is also a possibility that Ca^2+^ binds directly to tubulin to change the microtubule structure. Although *in vitro* microtubules are known to depolymerize in the presence of Ca^2+^, haptonematal MAPs may prevent the microtubules from disassembly. The electron microscopy images showed haptonema microtubules to be covered with associated proteins (Fig. 4D), which probably include more than one kind of MAP. These MAPs may stabilize the microtubules and prevent them from breakage even in the presence of Ca^2+^ or under mechanical strain.

The taxol-induced twist is converted into bending, which propagates from the distal to the proximal region of the haptonema by the addition of Ca^2+^, indicating that another mechanical state of haptonematal microtubules is induced by Ca^2+^. The direction of propagation of taxol-induced bending coincides with that of normal haptonematal coiling, supporting the idea that the Ca^2+^-dependent configuration change is linked to coiling.

In this study we observed structures associated with haptonematal microtubules (Fig. 4). In addition to the MAPs that cover the microtubule surface, we saw a filamentous structure that spread out from the bundle of haptonematal microtubules. This filament may be present with haptonematal microtubules to bundle them and might be involved in the Ca^2+^-dependent configuration change of microtubules for coiling. Another structure we observed was a mass of small particles at the tip of a haptonema, which may play roles both in fixing the distal end of the haptonema and in initiating structural changes of the haptonema for coiling, although it is still possible that these structures are a mass of depolymerized tubulins. Centrin is localized to haptonemata in a spotted pattern, mostly near the distal end (Lechtreck, 2004). However, it is not clear whether the mass of small particles that we observed (Fig. 4F) is simply comprised by centrins. Identification of the Ca^2+^-binding proteins will be key to understanding of the molecular mechanism of haptonematal coiling.

### Possible models for haptonematal coiling

To explain the mechanism of coiling, we present three models. In the first model, coiling is accounted for by Ca^2+^-dependent structural changes of MAPs that connect the neighboring microtubules (Fig. 8A). We hypothesize that the configuration of microtubules in the coiled haptonema represent a stable state when Ca^2+^ is bound to the MAPs (Fig. 7; Fig. 8A, right). In the absence of Ca^2+^, haptonematal microtubules are aligned approximately in parallel (Fig. 8A, left). When Ca^2+^ binds to the MAPs, their structural change imposes a tension between the neighboring microtubules, making the energy state of the haptonema higher, to induce twisting of the microtubules (Fig. 8A, middle), resulting in writhing to make a coil.

The second model (Fig. 8B) is based on our electron microscopy observation that one or two of the microtubules wind around the parallel bundle of the other microtubules (Fig. 4B; Fig. 8B, left). This model assumes that one of the seven haptonematal microtubules has specific Ca^2+^-biding MAPs that link this microtubule to the other six microtubules. Ca^2+^-binding to the MAPs may then change the structure of this special microtubule, enforcing the other six microtubules to twist (Fig. 8B, middle). The torsional force would be relaxed when the haptonema is coiled (Fig. 8B, right). This model is somewhat similar to the model that explains coiling of the *Vorticella* stalk, where a thin filament of a contractile protein, spasmin, winds around the elastic rod of the stalk. Once Ca^2+^ binds to spasmin, the thin filament quickly changes the conformation to coil the thick filament (Misra et al, 2010). This model agrees with the fact that one microtubule deviates from the other six microtubules in its distribution (Fig. 3G; Fig. S2). Therefore, it is reasonable to assume that one of the haptonematal microtubules is associated with specific Ca^2+^-binding proteins and drives the rapid coiling.

**Fig. 8.**
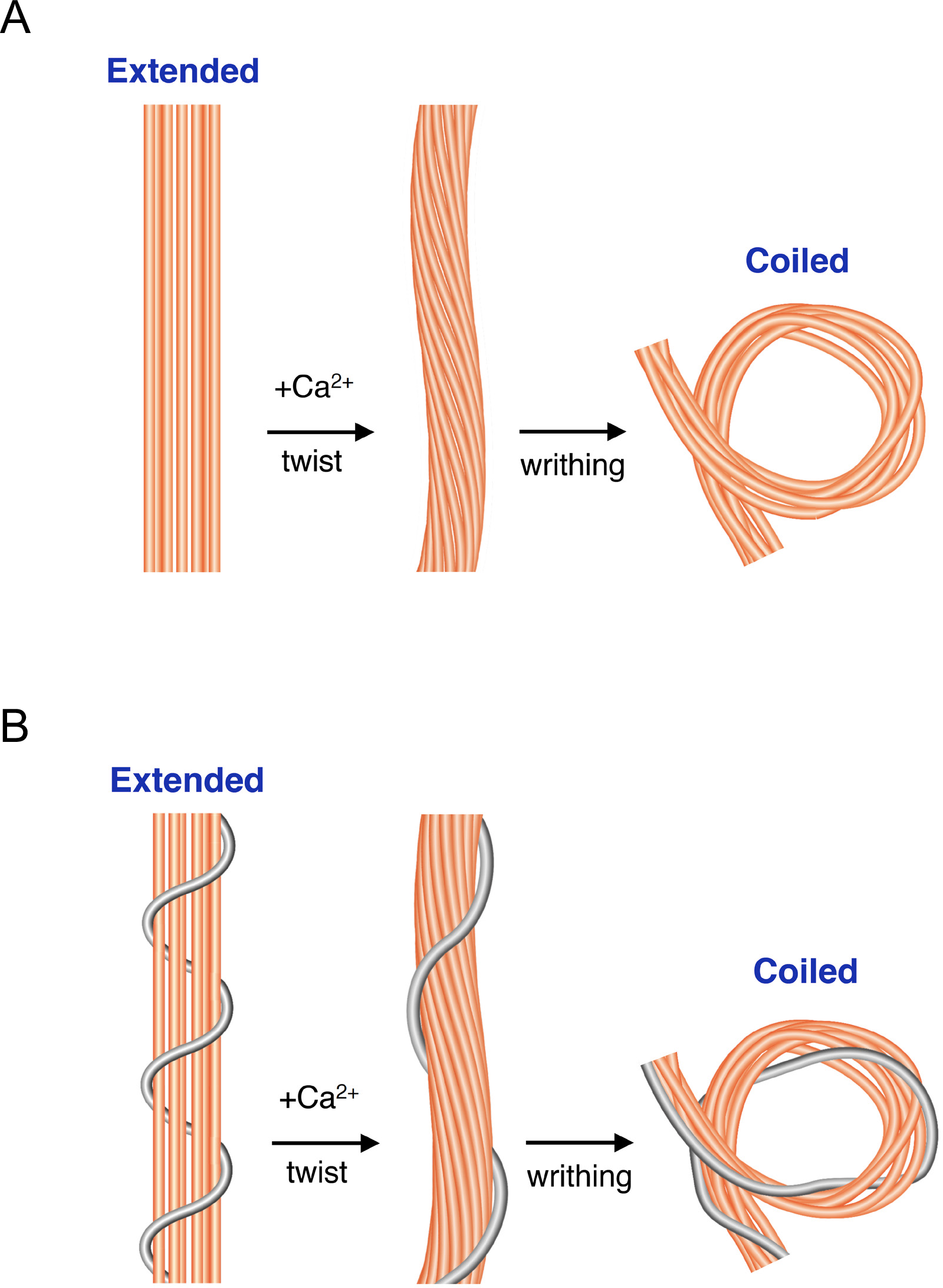
Two possible models for the mechanism of haptonematal coiling based on the twisting and writhing of haptonematal microtubules. (**A**) Microtubules in the extended haptonema are mostly parallel to each other in the absence of Ca^2+^ (left). Upon Ca^2+^ binding to MAPs that link the neighboring microtubules, the microtubules become twisted, which induces the formation of a gentle helix (middle). The haptonema would then form a coil by writhing, in which the microtubules are again mostly parallel, except that two opposite regions are twisted 2+ (right). This state is energetically stable with the aid of MAPs and Ca^2+^. (**B**) In the extended state, one microtubule winds around the other six haptonematal microtubules (left), connected with Ca^2+^-binding MAPs. Binding of Ca^2+^ induces the configuration change of this microtubule to twist the other microtubules, resulting in the formation of a gentle helix (middle). After writhing, the haptonema finally forms a coil (right).

The above two models attribute coiling of a haptonema to structural changes of the MAPs that link two haptonematal microtubules; however, the third model (Fig. S7) assumes association of Ca^2+^-biding MAPs to each protofilament that constitutes a microtubule. Structural changes of these MAPs would alter the conformation of the protofilament and its interaction with neighboring protofilaments (Fig. 7). These specific MAPs could bind either to one or two of the protofilaments (Fig. S7D), or to all of the protofilaments (Fig. S7G). The former case resembles the model explaining coiling of the bacterial flagellar filament, a helical machinery consisting of 11 protofilaments. They are composed of flagellin, which can adopt two slightly different conformations. The flagellar filament is straight when all the subunits have the same conformation but becomes helical when some of the protofilaments adopt the different structural state. In that way, the flagella can instantly switch from left-handed to right-handed helical structures (Calladine, 1978; Namba and Vonderviszt, 1998). In the case of a haptonema, if some of the MAPs specifically bind to one or two of the protofilaments, as is the case with axonemal doublet microtubules, these protofilaments would change their structure (Fig. S7D), resulting in coiling of the haptonematal microtubules (Fig. S7E). Coiling of the haptonema can be explained if the coiling of individual microtubules is coordinated (Fig. S7F).

Microtubule coiling could also occur when all of the protofilaments change their structure because of conformational change of the MAPs. Changes in the inter-protofilament interactions could twist the protofilaments (Fig. S7G), causing a torsional force on the microtubule, which may be relaxed when the microtubule coils (Fig. S7H, I). In all of these models, coiling of the haptonema is explained by conformational changes of the protofilaments or microtubules, which are made possible by Ca^2+^-dependent structural changes of MAPs associated with the protofilaments or microtubules (Fig. 7).

Microtubules are the most rigid cytoskeletal elements but can form flexible arcs with short wavelengths in living cells (Brangwynne et al, 2006). Some MAPs are known to cause conformational changes in protofilaments and overall microtubule structures, as seen in the case of stathmin (Sobel et al, 1989; Brouhard and Rice, 2014). The combination of taxol and MAPs has novel effects on the mechanical properties of microtubules (Hawkins et al., 2013), which is consistent with our present data. Further studies on the structures and Ca^2+^-dependent configurational changes of haptonematal microtubules and their associated proteins using cryoelectron microscopy and tomography should shed light on the mechanism of rapid coiling induced without microtubule-based motors. Identification of the Ca^2+^-binding MAPs will greatly clarify the molecular basis for the mechanism.

### Materials and methods

#### Haptophytes

*Chrysochromulina* sp. was isolated from Tokyo bay in 2013 using micropipette isolation (Anderson and Kawachi, 2005). It was then characterized (Fig. S1) and registered as a culture collection (NIES-4122). *Chrysochromulina* sp. NIES-4122 was maintained in Daigo IMK medium (Nippon Seiyaku Co., Osaka, Japan) at 20°C under a 14:10 h light: dark regime. The haptophyte could be recovered as a pellet after centrifugation but the haptonemata were mostly detached owing to the mechanical stimulus of colliding with the bottom of the centrifuge tube. To increase the number of cells in a microscopic field, we concentrated haptophytes using a non-toxic density gradient medium, Percoll. Percoll (GE healthcare) was diluted with two times concentrated artificial seawater (ASW; 460.3 mM NaCl, 10.11 mM KCl, 9.18 mM CaCl_2_, 35.91 mM MgCl_2_, 17.49 mM MgSO_4_, 0.1 mM EDTA and 10 mM HEPES-NaOH, pH 8.2) to make 50% Percoll. Ten milliliters of culture was layered on 100 μl of 50% Percoll in a 15 ml Falcon tube and was centrifuged at 2,800 g for 2 min. A hundred microliters of concentrated cells were collected from the boundary between the culture medium and Percoll.

#### Light microscopy observation of haptonematal coiling and uncoiling

Coiling and uncoiling were induced by tapping of the microscopic stage (Kawachi and Inouye 1994). Haptonemata were observed with a phase contrast microscope (BX51, Olympus, Tokyo, Japan). To record haptonematal coiling, the microscope was connected to a high-speed CCD HAS-D3 camera (Ditect, Tokyo, Japan). Haptonematal bending was analyzed from high-speed camera images using Bohboh software (Bohboh Soft, Tokyo, Japan). To examine the effects of Ca^2+^, 100 μl of concentrated cells were mixed with 900 μl of Ca^2+^-free ASW (462.01 mM NaCl, 9.39 mM KCl, 59.08 mM MgCl_2_, 10 mM HEPES, pH 8.0), 10 mM EGTA in Ca^2+^-free ASW, or 10 μM EGTA and 50 μM BAPTA-AM (Dojindo, 50 mM stock solution in DMSO) in Ca^2+^-free ASW. Paclitaxel (taxol) and nocodazole were dissolved in dimethyl sulfoxide (DMSO) to 20 mM and added to media to a final concentration of 20 μM.

#### Electron microscopy

Thin-section electron microscopy was performed as described previously (Konno et al, 2010) with some modifications. To examine microtubule configurations in sequential images, a flat surface of polymerized Epon resin was treated with 1 mg/ ml poly-lysine for 30 min, followed by coating with 10 mg/ml bovine serum albumin (BSA). Harvested haptophytes were fixed with 2.5% glutaraldehyde and were then deposited onto the poly-lysine coated surface by centrifugation at 200 × g for 10 min. The samples were post-fixed with 1% OsO_4_, dehydrated and embedded again according to a general procedure for thin-section electron microscopy. For negative staining, harvested cells were incubated in Ca^2+^-free ASW containing 10 mM EGTA for 10 min. Cells were then gently mounted on a carbon-coated Cu grid and demembranated with 0.5% NP-40 in 100 mM HEPES-NaOH (pH 7.2), 2 mM MgSO_4_, 5 mM EGTA for 15 sec. The grid was rinsed with the HEPES buffer for 1 min and negatively stained with 1% uranyl acetate. Electron microscopy was carried out using a JEOL 1010 or Tecnai F20 transmission electron microscope. Persistence lengths (*L_p_*) were calculated according to Gittes et al (1993) using the following equation:

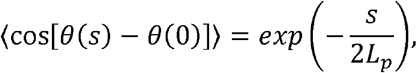

where *θ*(0) and *θ*(*s*) are the tangent angles of the filament at the origin (0) and at each point *s* along the filament. *s* denotes the arc length. Negative stain images were processed using ImageJ to obtain coordinates for calculation.

## Acknowledgements

We thank Shigekatsu Suzuki and Azumi Fukuda (University of Tsukuba) for their advice on haptophyte culture and Kangmin Yan (National Institute of Advanced Industrial Science and Technology) for her technical help with electron microscopy. We thank Jeremy Allen, PhD, from Edanz Group (www.edanzediting.com/ac) for editing a draft of this manuscript.

## Author contributions

K.In. conceived the study; K.A., T.N., K.Is. collected and identified haptophytes; M.N., K.A., K.H., K.S. K.In. designed experiments; M.N., K.A., K.H., K.S., K.In. performed experiments; N.M., K.A., K.S. analyzed data; K.I. wrote the manuscript; all authors agreed to the final version of the manuscript.

## Supplementary information

Supplementary figures and movies accompany this paper.

## Supplementary Figures

**Supplementary Fig. S1. Characterization of morphology and haptonematal coiling in *Chrysochromulina* sp. NIES-4122.**
(**A**) Negative staining image showing extracellular scales without spines. Bar, 100 nm. (**B**) A differential interference contrast image of a whole haptophyte. Bar, 10 μm. (**C**) Thin section electron microscopy image of the cell body. Bar, 1 μm. (**D**) Ca^2+^-dependent haptonematal coiling. Haptophytes were suspended in artificial sea water (ASW), Ca^2+^-free sea water (CFSW), CFSW with 10 mM EGTA, or CFSW with 10 mM EGTA plus 50 μM BAPTA-AM. Error bars represent the standard deviation. N=5. *P<0.01 (Student’s test).

**Supplementary Fig. S2. Changes in the center-to-center distance between adjacent haptonematal microtubules.** The center-to-center distances between adjacent microtubules were measured from six sequential thin-section electron microscopy images. (**A, B**) Sequential thin-section images of an extended (A) and coiled (B) haptonemata. The section numbers and the letters on the microtubules correspond to those in C and D. Bars, 100 nm. (**C, D**) Center-to-center distances between adjacent microtubules were plotted against the sequential section number, showing changes in the distance along the longitudinal axis of the haptonema. The extended haptonema (**C**) shows relatively constant distances among sequential sections but the coiled haptonema sometimes shows deviation for one of the inter-microtubule distances, which changes with the section number (**D**).

**Supplementary Fig. S3. Measurement of persistence length.**
(**A**) Plots along a negative stain image of a haptonema using ImageJ. The inset shows a magnified image of a part of the coiled region. (**B, C**) Distribution of persistence lengths. Green and red bars represent counts of the persistence length of the coiled and extended regions, respectively. The histogram in (B) is plotted with a bin width of 10 μm. (C) shows the region shorter than 10 μm, with the bin width of 1 μm.

**Supplementary Fig. S4. Definition of curvatures.**
(**A**) The curvature of a certain region of a haptonema was defined as the reciprocal of the radius of the inscribed circle (dashed blue or orange line). (**B**) Plot of curvature against distance from the base of a haptonema. The curvature values are either positive or negative, depending on the bending direction. The absolute value of curvature is used in Fig. 5F and Fig.S5 to show the extent of bending.

**Supplementary Fig. S5. Comparison of the maximum curvature of bends induced by microtubule-modifying drugs.**
Based on the analysis in Fig. 5, the average of absolute curvature values was calculated. Bars represent the standard deviation. N=20.

**Supplementary Fig. S6. Bend propagation of taxol-treated haptonemata.**
The curvature along the haptonema was measured at 5, 30 and 60 min after treatment with taxol. (**A**) In the presence of Ca^2+^ (ASW); (**B**) In the absence of Ca^2+^ (CFSW+EGTA+BAPTA-AM). Note that the peak of the maximum curvature (arrows) propagates toward the base of the haptonema with time in the presence of Ca^2+^ and that the gentle helix propagates toward the tip of the haptonema with time in the absence of Ca^2+^.

**Supplementary Fig. S7. Basic model for the microtubule coiling mechanism.**
Diagrams illustrating the arrangement of tubulin dimers in a microtubule (**A, D, G**), the state of a microtubule (**B, E, H**), and the state of a bundle of seven microtubules as in a haptonema (**C, F, I**). The Ca^2+^-bound MAPs that are responsible for coiling are associated with either a set of protofilaments (D-F) or to all protofilaments (G-I). When Ca^2+^ is not bound to the MAPs, the microtubules are straight in both cases (A-C). In **D**, Ca^2+^-binding to the MAPs changes the structure of the protofilament to which the MAPs are attached (pink protofilament). If we assume a mechanism similar to that of bacterial flagella, a slight decrease in the subunit spacing and changes in the interaction between neighboring protofilaments result in bending and twisting of the microtubule (D), which lead to coiling of the microtubule (E) and thus coiling of the haptonema (F). In **G**, Ca^2+^ binding to the MAPs alters the structures of all of the protofilaments and inter-protofilament interactions. This would twist the protofilaments, imposing a torsional force on the microtubule, and make the coiled conformation more stable.

**Movie 1. Real-time movie of haptonematal coiling and uncoiling.**
The coiling and uncoiling were induced by tapping the microscopic stage. The movie was recorded at 100 fps and plays at real time. The process of coiling is too rapid to be seen but that of uncoiling proceeds more slowly.

**Movie 2. High-speed movie of haptonematal coiling.**
The movie was recorded at 1000 fps and plays at a speed of 0.03×.

**Movie 3. High-speed movie of haptonematal uncoiling.**
The movie was recorded at 200 fps and plays at a speed of 0.15×.

